# Mitochondrial dysfunction heightens the integrated stress response to drive ALS pathogenesis

**DOI:** 10.1101/2024.05.13.594000

**Authors:** Curran Landry, James Costanzo, Miguel Mitne-Neto, Mayana Zatz, Ashleigh Schaffer, Maria Hatzoglou, Alysson Muotri, Helen Cristina Miranda

## Abstract

Vesicle-associated membrane protein-associated protein-B (VAPB) is an ER membrane bound protein. VAPB P56S causes a dominant, familial form of amyotrophic lateral sclerosis (ALS), however, the mechanism through which this mutation causes motor neuron (MN) disease remains unknown. Using inducible wild type (WT) and VAPB P56S expressing iPSC-derived MNs we show that VAPB P56S, but not WT, protein decreased neuronal firing and mitochondrial-ER contact (MERC) with an associated age-dependent decrease in mitochondrial membrane potential (MMP); all typical characteristics of MN-disease. We further show that VAPB P56S expressing iPSC-derived MNs have enhanced age-dependent sensitivity to ER stress. We identified elevated expression of the master regulator of the Integrated Stress Response (ISR) marker ATF4 and decreased protein synthesis in the VAPB P56S iPSC-derived MNs. Chemical inhibition of ISR with the compound, ISRIB, rescued all MN disease phenotype in VAPB P56S MNs. Thus, our results not only support ISR inhibition as a potential therapeutic target for ALS patients, but also provides evidence to pathogenesis.

## INTRODUCTION

Amyotrophic Lateral Sclerosis (ALS), also known as Lou Gehrig’s disease, is the most common adult–onset motor neuron disease. There is currently no cure for this devastating disease that affects approximately 30,000 individuals at any time in United States.^1^ A novel, autosomal dominant form of ALS was mapped to chromosome 20q13.3 in a Brazilian family in 2004.^2^ Subsequently, the mutation (NM_004738.5:c.166C>T, p.(Pro56Ser); hereafter referred to as P56S) in the *Vesicle Associated Membrane Protein (VAMP) Associated Protein B (VAPB)* gene was identified.^2,3^ VAPB contains three different domains, the transmembrane domain, the coiled-coiled domain and the major sperm protein (MSP) domain containing the double phenylalanine in an acidic tract (FFAT) binding motif, and the MSP domain is highly conserved across multiple species including humans, *D. melanogaster, S. cerevisiae*.^4^ VAPB is an ER membrane bound protein that tethers binding proteins and organelles to the ER, including the mitochondria, the Golgi complex, and the plasma membrane.^5,6^ The MSP domain is known for its protein binding ability and the VAPB P56S mutation, located within the MSP domain, impairs VAPB’s tethering function.^3,5,7,8^

One of the main organelles that VAPB anchors to the ER is the mitochondria. VAPB has been shown to tether the mitochondria to the ER through binding of the mitochondrial tethering protein, protein tyrosine phosphatase interacting protein-51 (PTPIP51), and this binding is disrupted by the VAPB P56S mutation.^9,10^ It has also been shown that this interaction is present at synapses, and disruptions to it affect synaptic activity.^11^ Furthermore, the P56S mutation disrupts VAPB normal binding to FFAT (double phenylalanine in an acidic tract) motif containing proteins, and is also known to cause a conformational change leading VAPB proteins to aggregate. Protein aggregates within cells expressing both wild type VAPB (VAPB WT) and VAPB P56S have been shown to include VAPB WT into the aggregates, demonstrating sequestration of VAPB WT as another possible mechanism for pathogenesis. ^12,13^ VAPB WT interactome studies have been previously performed to help elucidate VAPB functions, however they have not been carried out in a disease context, in comparison to mutant VAPB P56S, or in motor neurons, an ALS affected cell type.^14,15^ A previously published VAPB iPSC-derived disease model showed ALS type VIII patient iPSC-derived neurons have decreased levels of soluble VAPB.^8^

In this study we generated iPSCs stably expressing either the VAPB P56S or VAPB WT in a CRISPR-Cas9 knockout VAPB background to interrogate VAPB P56S specific pathogenesis associated with ALS. We differentiated the iPSC into motor neurons and identified an electrophysiological activity dysregulation by a multi-electrode array (MEA) system. We then explored the molecular mechanism of this dysregulation and found that VAPB P56S has reduced binding with many mitochondrial binding partners, including PTPIP51.^9,16^ Given this, we examined the mitochondrial-ER contacts (MERC) and found that the VAPB P56S motor neurons displayed decreased MERC compared to VAPB WT throughout differentiation.

Moreover, we identified an age-dependent decline in mitochondrial membrane potential (MMP). Importantly, we observed that the VAPB P56S iPSC-derived motor neurons were more sensitive to cell stressors, exhibiting early activation of the Integrated Stress Response (ISR), namely through the DELE1 mediated pathway triggered by mitochondrial stress. Using an ISR inhibitor (ISRIB) we were able to rescue the neuronal firing and MMP dysfunctions, indicating that they were manifestations of ISR activation. To our knowledge, this is the first systematic investigation of VAPB WT and VAPB P56S in ALS disease relevant cells, iPSC-derived motor neurons of the same genetic background, elucidating the most complete pathogenesis of ALS type VIII to date.

## RESULTS

### VAPB P56S Motor Neurons Display Decreased Neuronal Firing

To create isogenic iPSC lines stably expressing either VAPB WT and VAPB P56S first we knocked out endogenous VAPB using CRISPR-Cas9 technology and then generated lentiviral expression vectors containing the mutant or the wild-type genes under the control of a tetracycline-dependent promoter (Extended Data Figure 1a-c). Once stably transduced and selected, VAPB expression is controlled in a dose-dependent manner with addition of doxycycline. A doxycycline dose-response curve for exogenous VAPB expression in the transduced iPSCs was performed in comparison to endogenous VAPB expression in patient iPSC harboring the VAPB P56S mutation and familial unaffected controls (VAPB WT) to select the physiological doxycycline dose for both VAPB WT and VAPB P56S (0.2 ug/mL) (Extended Data Figure 2a). All experiments were performed in iPSC-derived motor neurons, differentiated using a dual SMAD inhibition protocol, as previously described (**Figure 1a**).^17^ The cultures predominantly contain mature motor neurons (MN) on day 25. iPSCs were treated with doxycycline to express either VAPB WT or VAPB P56S from Day 1 of differentiation to motor neurons. No significant difference was noted between the VAPB WT and VAPB P56S doxycycline inducible lines, in either efficiency of motor neuron progenitor (Extended Data Figure 2b) or motor neuron differentiation (Extended Data Figure 2c).

**Figure 1.**
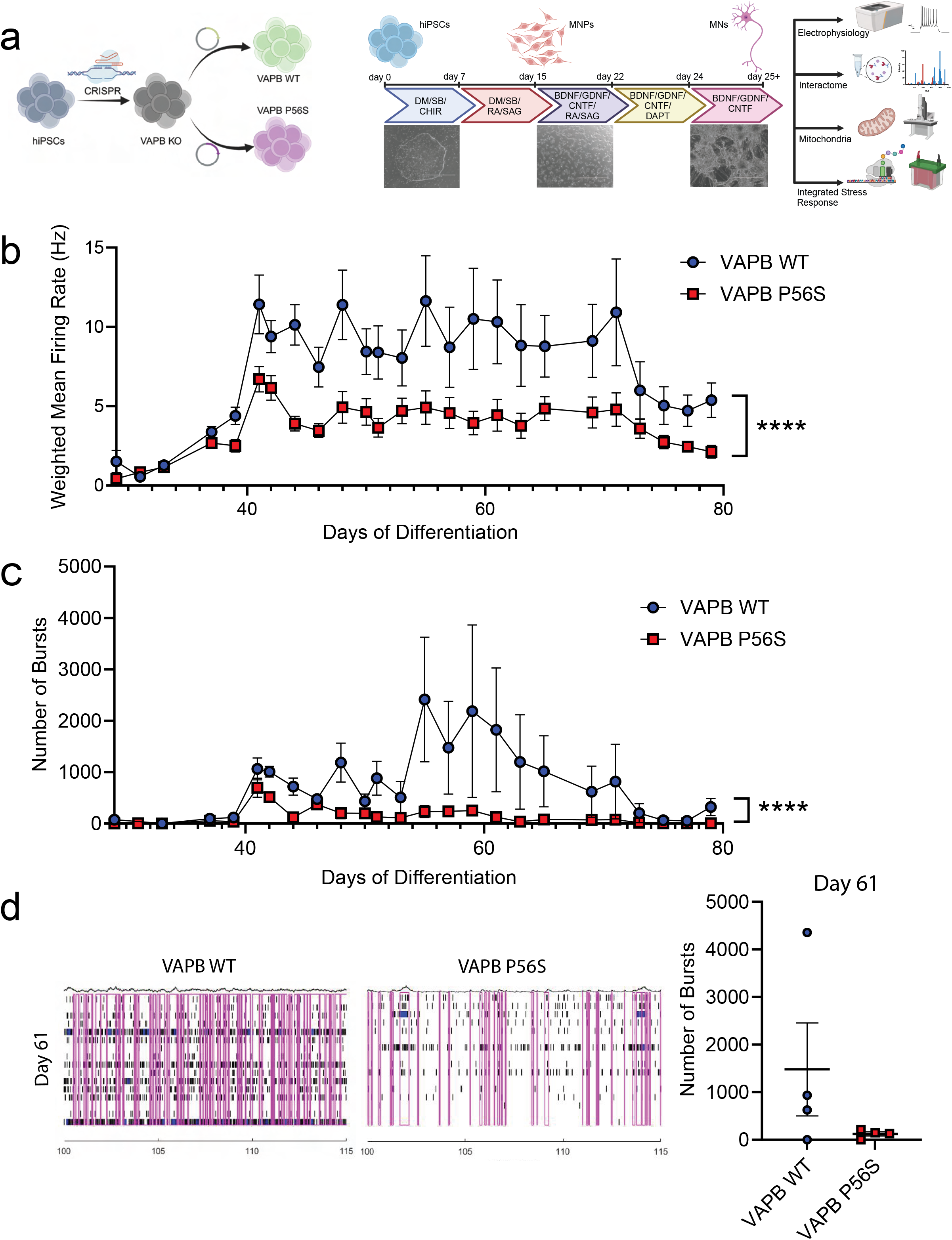
VAPB P56S motor neurons exhibit decreased neuronal firing rate compared to WT controls. a) Schematic depicting the timeline of factors used in the motor neuron differentiation protocol. Made with Biorender.com. b) Weighted mean firing rate through day 80 of motor neuron differentiation. Mean ± SEM, N=8 separate wells of an MEA until day 55, at which they are reduced to four wells, mixed-effects analysis (REML), interaction effect *p*<0.0001**** c) Number of bursts through day 80 of motor neuron differentiation. Mean ± SEM, N=8 separate wells of an MEA until day 55, at which they are reduced to four wells, mixed-effects analysis (REML) interaction effect *p*<0.0001****d) Raster plot of neuronal firing on day 61, 100 seconds after starting recording, bursts outlined in magenta. Scatter plot quantifying total number of bursts per well on day 61 of differentiation.

**Figure 2.**
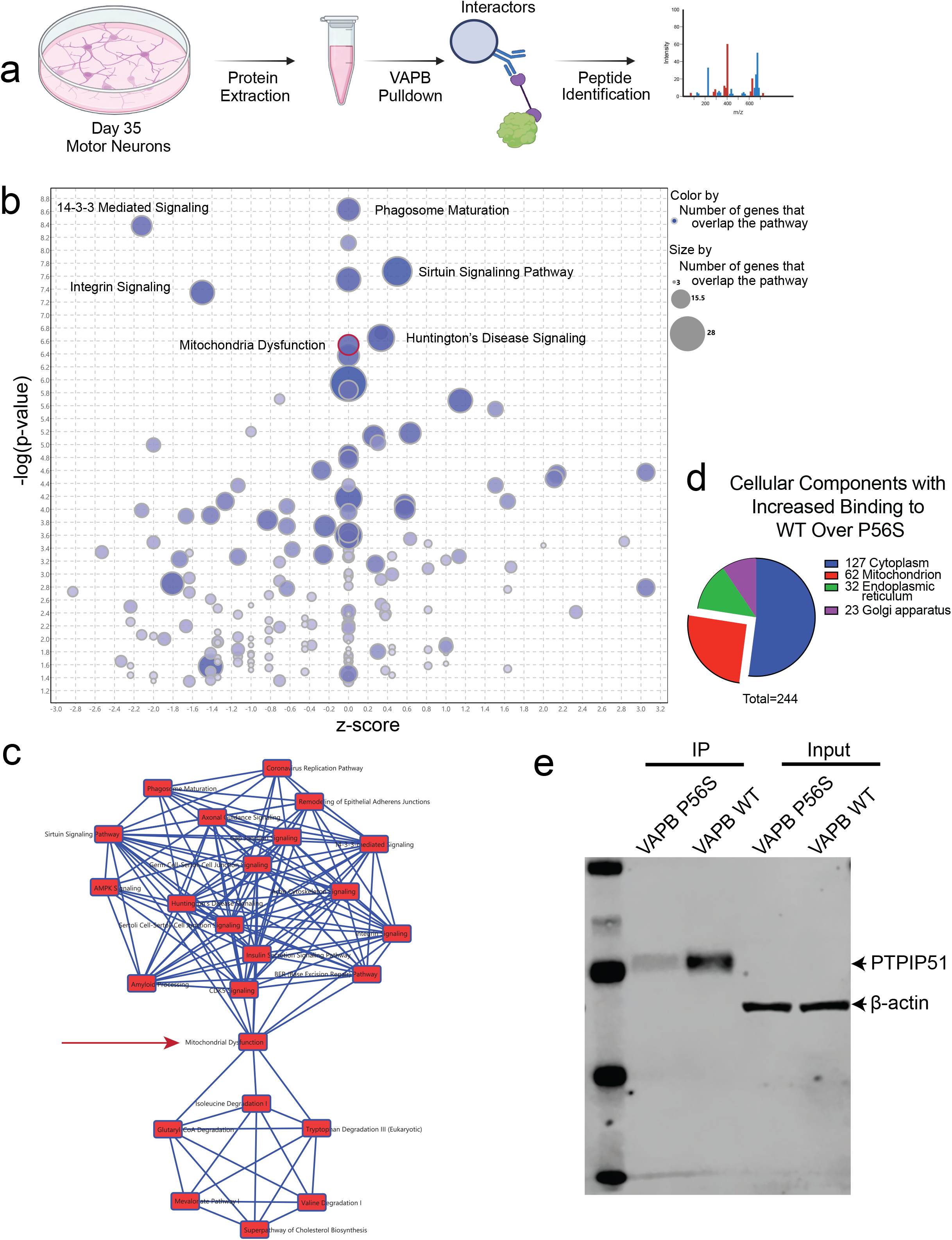
Interactome Analysis of VAPB WT and VAPB P56S. a) Schematic depicting the coimmunoprecipitation workflow to isolate and identify VAPB interactors. b) IPA analysis depicting pathways highly enriched with proteins that have increased binding to WT over P56S. The z-score (X-axis) is IPA’s prediction of activation or inhibition of the pathway, while the Y-axis is -log(p-value) for predicted significance of pathway enrichment. Size of the circle indicates raw number of proteins in a given pathway found in our dataset, while shading depicts percentage of proteins present in both the canonical pathway and our dataset. c) Diagram depicting connections between highly enriched pathways in IPA. d) Cellular compartment analysis using DAVID of proteins with 1.5-fold increase in binding of VAPB WT over VAPB P56S. e) Western Blot of VAPB IP samples and corresponding input stained for PTPIP51 and ß-actin.

To examine whether the VAPB P56S mutation affects neuronal activity, a clinically relevant phenotype to ALS, we used a multielectrode array (MEA) system. Mature VAPB P56S and VAPB WT iPSC-derived motor neuron recordings were taken every other day and compared. Just after day 40, the iPSC-derived motor neuron cultures reached a plateau in electrophysiological activity. The VAPB P56S iPSC-derived motor neurons show significantly decreased firing from day 40 onward when compared to VAPB WT control (**Figure 1b**).

Additionally, we quantified the neuronal bursting of our iPSC-derived motor neurons to assess whether the P56S mutation was affecting the firing rate independently of bursting or showing a commensurate decrease in the network firing of the VAPB P56S iPSC-derived motor neurons compared to VAPB WT controls. We found that only the VAPB WT iPSC-derived motor neurons show consistently increased bursting after day 55 (**Figure 1c**). We also examined histogram and scatter plots of the neuronal activity later in differentiation (day 61) to assess the specific firing of the aged neurons, as ALS is a disease that presents with age (**Figure 1d**). We saw that the VAPB WT iPSC-derived motor neurons had increased bursting in addition to their increased firing rate compared to the P56S neurons, indicating greater connectivity and synchronicity within the VAPB WT iPSC-derived motor neuronal cultures.

### VAPB P56S leads to loss of Mitochondrial binding partners

VAPB is a tethering protein, therefore when we began to search for a molecular pathogenesis we decided to analyze the binding partners of VAPB and how the P56S mutation affects them. Our model system allowed us to isolate VAPB P56S binding partners and define the VAPB P56S interactome independently of VAPB WT, something unachievable in patient lines as all known patients are heterozygous for the P56S mutation. To this end we cultured VAPB WT expressing, and VAPB P56S expressing, and VAPB KO iPSC-derived motor neurons and isolated VAPB associated proteins from them on day 35. The immunoprecipitated proteins were then submitted for mass spectrometry-based peptide identification (**Figure 2a**).

We compared the VAPB WT and VAPB P56S associated proteins to identify bindings that were disrupted in the VAPB P56S interactome. The list of differentially bound interactors was then used with Ingenuity Pathway Analysis (IPA) to reveal pathways in which the binding of VAPB would be most disrupted under disease conditions (**Figure 2b)**. We then had IPA generate connections between the most highly enriched pathways and found two clusters whose only connection between them was the pathway of mitochondrial dysfunction (**Figure 2c**). Given the possible significance of the mitochondrial dysfunction pathway identified in IPA, we also used the Database for Annotation, Visualization and Integrated Discovery (DAVID) to identify cellular compartments enriched for proteins with loss of binding with VAPB P56S compared to VAPB WT (**Figure 2d**). The compartment with the second highest number of interactors lost is the mitochondria, succeeding the cytoplasm, the largest cellular compartment. The results from both IPA and DAVID implicated mitochondrial binding partners as the disrupted interaction in VAPB P56S expressing iPSC-derived MNs corroborates previously published data showing that VAPB is a critical component in tethering the mitochondria to the ER, through an interaction with PTPIP51, an interaction known to be disrupted by the VAPB P56S mutation and results in decreased MERC.^9,10,16^ We confirmed the co-immunoprecipitation and mass spectrometry results through Western Blot, showing that PTPIP51 was enriched in the immunoprecipitation as well as showed decreased binding to VAPB in the VAPB P56S iPSC-derived motor neurons (**Figure 2e** and Extended Data Figure 3a).

**Figure 3.**
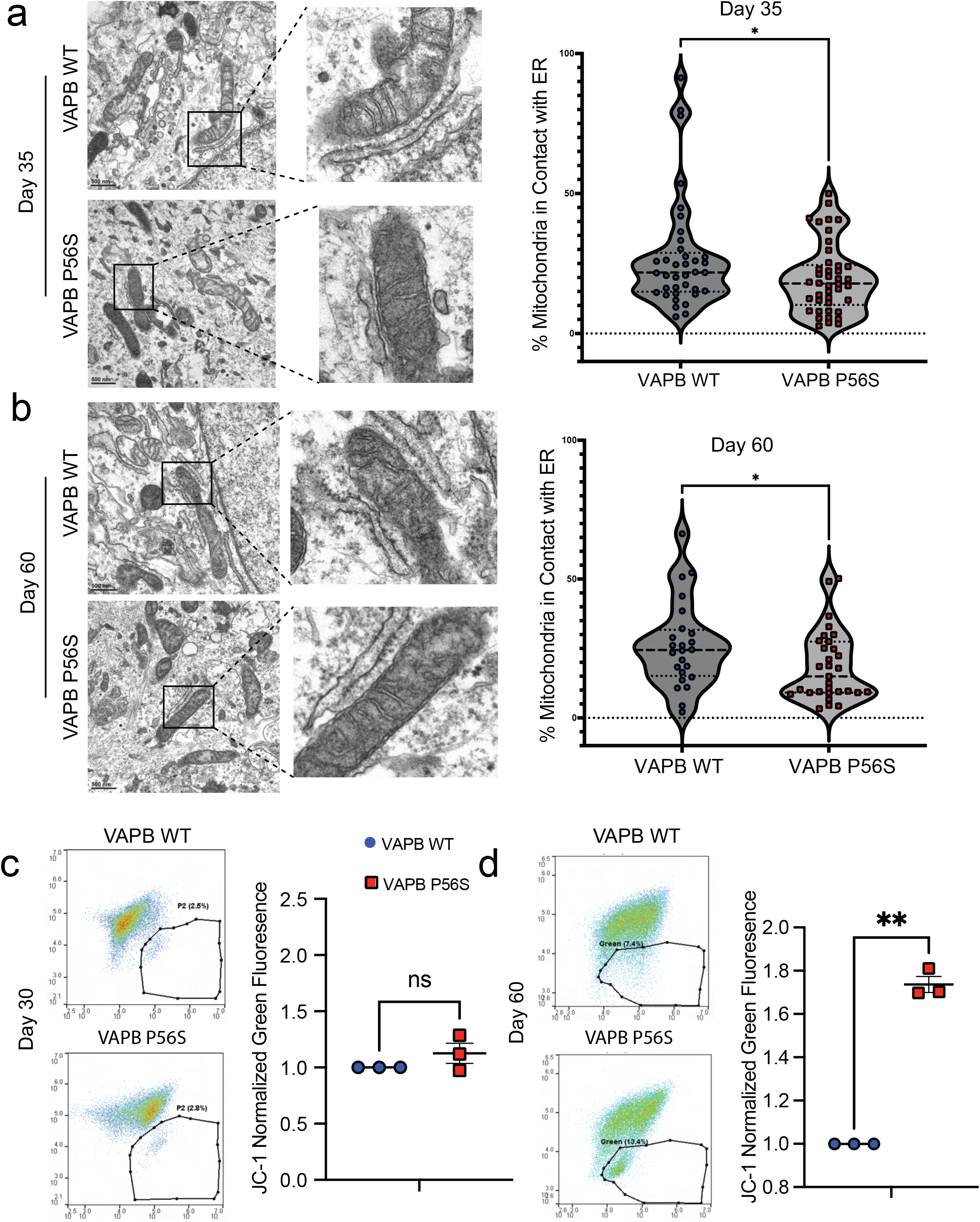
VAPB P56S Causes Reduced ER-Mitochondrial Contact and Decreased Mitochondrial Membrane Potential. a) Electron microscopy images taken of day 35 VAPB WT and VAPB P56S motor neurons with inset zooms on mitochondria with ER contact. Violin plot with median, first and third quartiles denoted as dashed lines, from samples submitted for EM, N=13-21 images analyzed for each condition, unpaired t-test *p*=0.0475* b) Electron microscopy images taken of day 60 VAPB WT and VAPB P56S motor neurons with inset zooms on mitochondria with ER contacts, Violin plot with median, first and third quartiles denoted as dashed lines, from samples submitted for EM, N=13-21 images analyzed for each condition, unpaired t-test *p*=0.0485* c) Scatter plot of day 30 motor neurons stained with JC-1 mitochondrial dye. The X-axis is red fluorescence, Y-axis is green fluorescence, gate is surrounding highly green fluorescing population. Mean ± SEM, N=3, paired t-test *p*>0.05. d) Scatter plot of day 60 motor neurons stained with JC-1 mitochondrial dye. X-axis is red fluorescence, Y-axis is green fluorescence, gate is surrounding highly green fluorescing population. Mean ± SEM, N=3, paired t-test *p*=0.0025**.

### The VAPB P56S mutation leads to a reduction in ER-mitochondrial contact and decreased mitochondrial membrane potential

To investigate if the loss of VABP P56S association to proteins in the mitochondria could alter the amount of MERC, we performed electron microscopy analysis to quantify the membrane contact sites between the ER and the mitochondria. We analyzed day 35 and day 60 VAPB WT expressing and VAPB P56S expressing iPSC-derived motor neurons, to examine not only if VAPB P56S affects MERC, but whether aging the iPSC-derived motor neurons may alter MERC, since we observed decreased neuronal firing after day 40 of differentiation (**Figure 1b**). We found that the VAPB P56S iPSC-derived motor neurons exhibited a significant decrease in the percentage of mitochondrial contact with the ER on both day 35 (**Figure 3a**) and day 60 (**Figure 3b**) of differentiation, corroborating previous reports (Extended Data Figure 4a-c).^9,10,18^ With this loss of MERC confirmed in our model, we next wanted to assess the functional consequence on the mitochondria. To investigate this, we used the JC-1 assay to measure the mitochondrial membrane potential (MMP). Since VAPB P56S iPSC-derived motor neurons display a decrease in electrophysiological firing after day 40, we decided to assess MMP throughout iPSC-derived motor neuron aging. Therefore, we assayed VAPB WT and VAPB P56S iPSC-derived motor neurons on days 30 and 60 of differentiation. We saw that the day 60 VAPB P56S iPSC-derived motor neurons had a significantly increased green fluorescing population, indicating decreased MMP compared to the VAPB WT controls, whereas on day 30, there was no significant difference between VAPB WT and VAPB P56S iPSC-derived motor neurons (**Figure 3c&d**). These findings suggest that although the reduction in MERC is reduced throughout the lifespan of the VAPB P56S iPSC-derived motor neurons, the functional impact on the MMP requires time to manifest.

**Figure 4.**
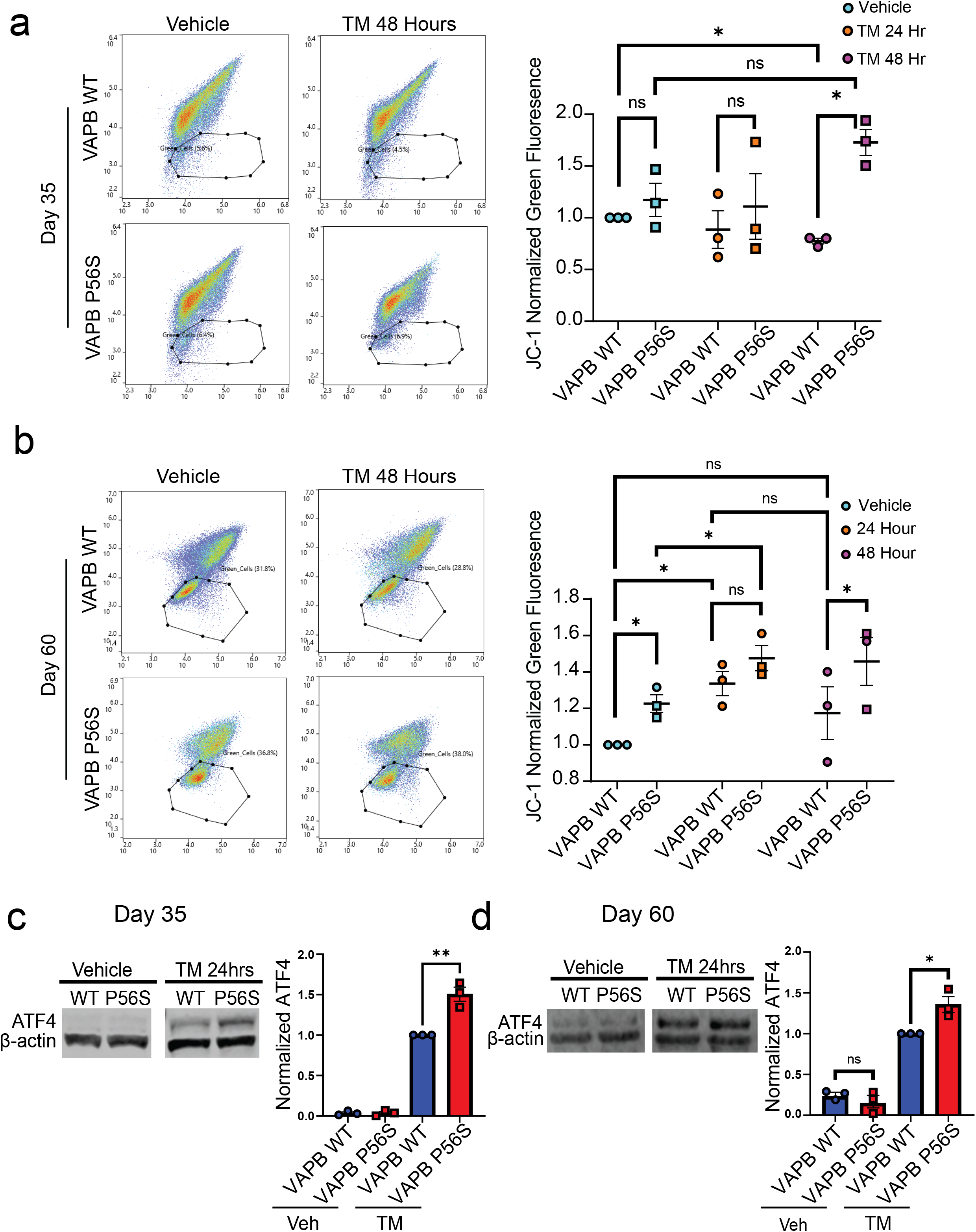
VAPB P56S Impairs Cellular Ability to React to Stress. a) Scatter plot of day 35 motor neurons stained with JC-1 mitochondrial dye after 24 hours, 48 hours or untreated with tunicamycin. Mean ± SEM, N=3, paired t-test VAPB WT Vehicle vs VAPB WT TM 48Hr *p*=0.0141*, VAPB WT TM 48Hr vs VAPB P56S TM 48Hr *p*=0.0450* b) Same as a, except on day 60 of differentiation. Mean ± SEM, N=3, paired t-test VAPB WT Vehicle vs VAPB P56S Vehicle *p*=0.043*, VAPB WT Vehicle vs VAPB WT TM 24Hr *p*=0.0374*, VAPB P56S Vehicle vs VAPB P56S TM 24Hr *p*=0.0229*, VAPB WT TM 48Hr vs VAPB P56S TM 48Hr *p*=0.0212* c) Western Blot of ATF4 basal levels and after 24 hours of tunicamycin exposure in day 35 motor neurons, Mean ± SEM, N=3, unpaired t-test *p*=0.0045** d) Same as c except on day 60 of differentiation. Mean ± SEM, N=3, unpaired t-test *p*=0.0217*

### VAPB P56S impairs the cellular ability to adapt to cell stressors

To test whether the physical disruption of the MERC is underlying the decreased MMP, the cells were subjected to stressors both early and late in differentiation. We hypothesized that if the decrease in MERC was underlying the decrease in MMP seen later in differentiation, inducing cell stress early in differentiation could mimic the decreased MMP. We selected tunicamycin, an ER stressor, as VAPB is an ER protein, and if the decreased MMP could be caused, at least partially, by loss of MERCs, ER stress would likely exacerbate it. To this end, we treated iPSC-derived motor neurons with tunicamycin (TM), for either 24 or 48 hours, and then assayed MMP. On day 35, there was no change in the MMP until 48 hours of treatment, after which the VAPB P56S iPSC-derived motor neurons display a significant decrease in MMP compared to the VAPB WT (**Figure 4a**). This indicates that VAPB P56S iPSC-derived motor neurons are more sensitive to stress compared to VAPB WT, with VAPB P56S iPSC-derived motor neurons exhibiting a significantly lower MMP compared with VAPB WT after 48 hours of TM dosing.

When repeating the same TM dosing experiment on day 60 of differentiation, a different pattern presents. After 24 hours of TM treatment, both VAPB WT and VAPB P56S iPSC-derived motor neurons have decreased MMP; however, after 48 hours, the VAPB WT have adapted to increase their MMP, and are no longer significantly different than the VAPB WT vehicle control. In contrast, the VAPB P56S iPSC-derived motor neurons show persistently decreased MMP at the 48-hour mark, indicating a failure to adapt to the TM challenge (**Figure 4b**).

In addition to examining the MMP of the cells after inducing stress at both timepoints, we also wanted to examine a common cellular stress response pathway marker We chose activating transcription factor 4 (ATF-4) as it is a central convergence point for several stress pathways, responsible for the enhanced transcription of many cellular stress-response proteins.^19-21^ We see no basal expression of ATF-4 in either VAPB WT or VAPB P56S iPSC-derived motor neurons on day 35, and only low-level expression on day 60, with no difference between VAPB WT and VAPB P56S. However, after TM treatment, both lines increase ATF-4 expression, however, VAPB P56S expresses a significantly higher level of ATF-4 than the VAPB WT in both day 35 and day 60 iPSC-derived motor neurons (**Figure 4c&d**).

Taken together these data suggest that not only is the physical disruption of the MERC underlying the functional deficits seen later in differentiation, but also suggests that the VAPB P56S iPSC-derived motor neurons have an increased sensitivity to stressors, and an impaired adaptation to them.

### VAPB P56S causes increased activation of the Integrated Stress Response paralleled by cleavage of mitochondrial protein DELE1

Once we established the increased sensitivity to stress of the VAPB P56S iPSC-derived motor neurons, we wanted to test whether they also exhibited molecular markers of mitochondrial stress in addition to the functional phenotype of decreased MMP we have seen later in differentiation. We decided to examine DELE1 as mitochondrial stress causes cleavage of DELE1 (also known as DELE1-L) into small (DELE1-s) fragments.^22-24^ When we assessed the levels of DELE1 we found day 35 VAPB P56S iPSC-derived motor neurons display increased levels of DELE1-s compared to VAPB WT controls with no difference in levels of DELE1-L (**Figure 5a**).

**Figure 5.**
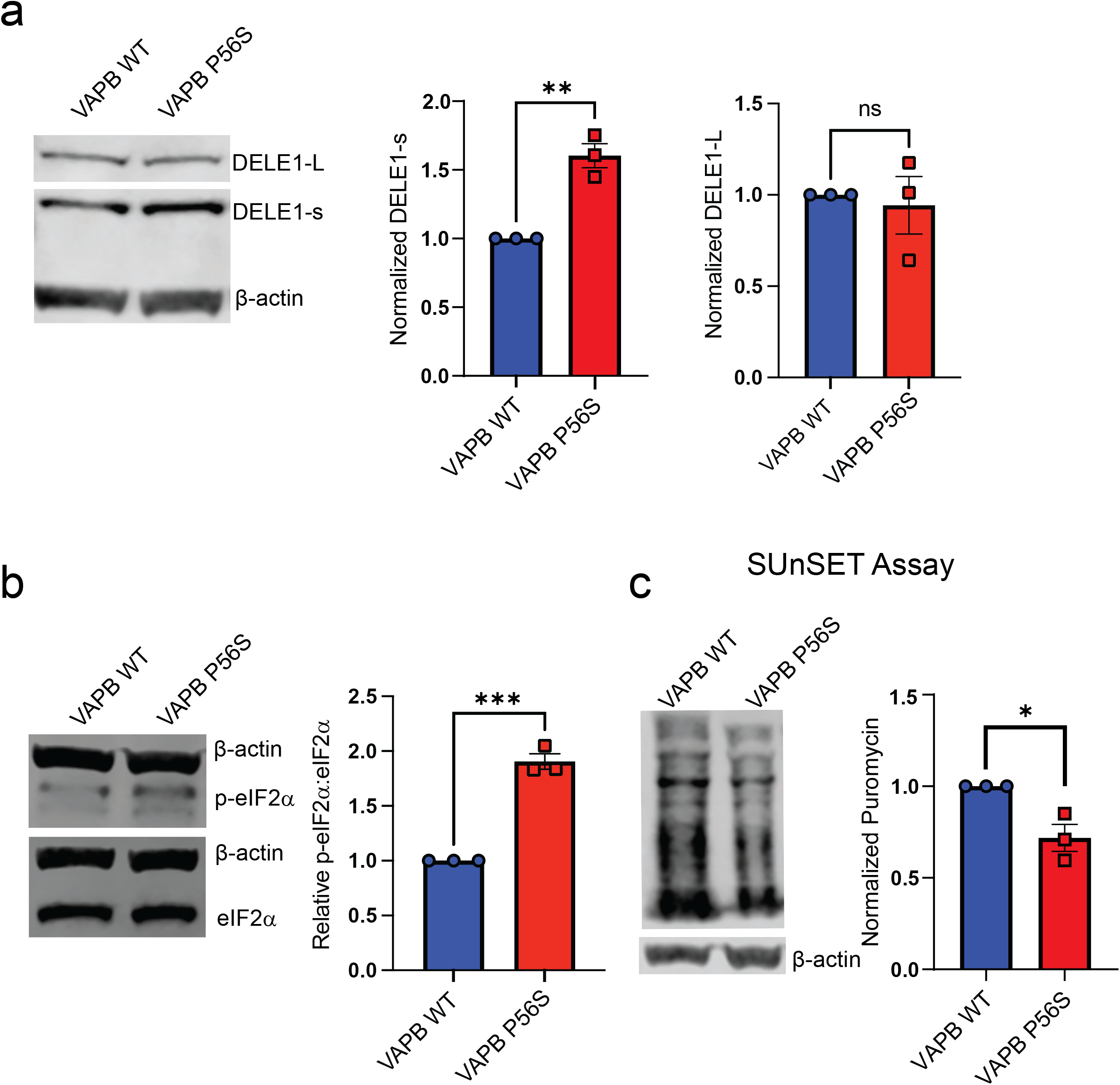
The Integrated Stress Response Pathway is activated early in VAPB P56S motor neuron differentiation. a) Western Blot for DELE1 large and small fragment on day 35 of differentiation Mean ± SEM, N=3, unpaired t-test *p*=0.0023** b) Western Blot for peIF2α and eIF2 on day 35 of differentiation. Mean ± SEM, N=3, unpaired t-test *p*=0.0002*** c) SUnSET assay on day 35 of differentiation. Mean ± SEM, N=3, unpaired t-test *p*=0.0188*

Previous work shows that mitochondrial stress leads to DELE1 cleavage, and once cleaved, DELE1-s then binds heme-regulated eIF2α kinase (HRI), activating it, leading to the phosphorylation of eIF2α.^22-24^ Once phosphorylated, p-eIF2α is known to cause expression of ATF-4, a key regulator in the Integrated Stress Response Pathway (ISR). In addition, the ISR is a key cellular process that is known to be linked to ER and mitochondrial health.^18,25,26^ Given this and our data showing disruptions in MERC, mitochondrial function, and ATF-4 response to stressors, we decided to examine the ISR in the VAPB P56S iPSC-derived motor neurons. We analyzed p-eIF2α as the downstream effector of the increase in DELE1-s and saw that the day 35 VAPB P56S iPSC-derived motor neurons exhibit a higher ratio of p-eIF2α:eIF2α than their VAPB WT counterparts (**Figure 5b)**.

p-eIF2α has two major canonical activities, activation of ATF-4 and global inhibition of mRNA translation, therefore we also examined the level of protein synthesis in day 35 motor neurons.^27,28^ To examine this, we utilized the Surface Sensing of Translation (SUnSET) assay to ascertain the level of translation through rate of puromycin incorporation.^29,30^ We observed a significant decrease in the amount of puromycin incorporation in the VAPB P56S iPSC-derived motor neurons, signaling a decrease in the level of translation on day 35, consistent with the elevated ratio of p-eIF2α:eIF2α (**Figure 5c**).

### Decreased MMP, neuronal firing, and protein synthesis in ALS type VIII iPSC-derived motor neurons can be rescued through inhibition of the ISR

Having demonstrated that the VAPB P56S mutation causes ISR activation in ALS type VIII iPSC-derived motor neurons, we sought to discern whether it caused the observed functional phenotypes. To do this, we treated iPSC-derived motor neuron cultures with Integrated Stress Response inhibitor (ISRIB), which when bound to eIF2B (a key component of a translational initiation complex) induces a conformational change in eIF2B prompting dissociation of p-eIF2α thereby promoting translation, inhibiting ATF4 expression, and blunting the ISR.^31^ We first examined ISRIB’s effect on protein synthesis using the SUnSET assay on day 35 and observed that ISRIB treatment significantly increased puromycin incorporation in VAPB P56S iPSC-derived motor neurons. Protein synthesis levels in ISRIB treated VAPB WT and VAPB P56S was comparable, indicating that ISRIB treatment restored decreased protein synthesis caused by VAPB P56S-induced ISR (**Figure 6a**). As expected, protein synthesis did not significantly change in VAPB WT expressing iPSC-derived motor neurons with ISRIB dosing, likely due to limited ISR activation in these cells. We wanted to next examine the effect of ISRIB on the decreased MMP observed in VAPB P56S expressing iPSC-derived motor neurons. After treatment with ISRIB for 24 hours on day 60, both VAPB WT and VAPB P56S iPSC-derived motor neurons showed a significant increase in MMP, with no significant difference between the two lines after treatment, indicating ISRIB does ameliorate the decreased MMP phenotype, and that the increased ISR activity in VAPB P56S iPSC-derived motor neurons is likely responsible for the decreased MMP (**Figure 6b**). Finally, we wanted to examine whether the decreased neuronal firing observed in VAPB P56S iPSC-derived motor neurons could be rescued through ISRIB treatment. Therefore, we began ISRIB dosing on day 45 of differentiation and observed a significant increase in the firing rate of VAPB P56S iPSC-derived motor neurons treated with ISRIB compared to that of vehicle treated VAPB P56S iPSC-derived motor neurons (**Figure 6c**). Along with decreased MMP and translation, the reduction in neuronal firing was at least partially caused by an activation of the ISR within the VAPB P56S iPSC-derived motor neurons. Taken together, our data suggests that the VAPB P56S mutation causes a disruption in PTPIP51 binding, leading to a decrease in MERC. This causes mitochondrial stress, which in turn activates the Integrated Stress Response Pathway through DELE1 cleavage. Activation of the ISR then leads to decreased neuronal firing and decreased MMP later in differentiation (**Figure 7**).

**Figure 6.**
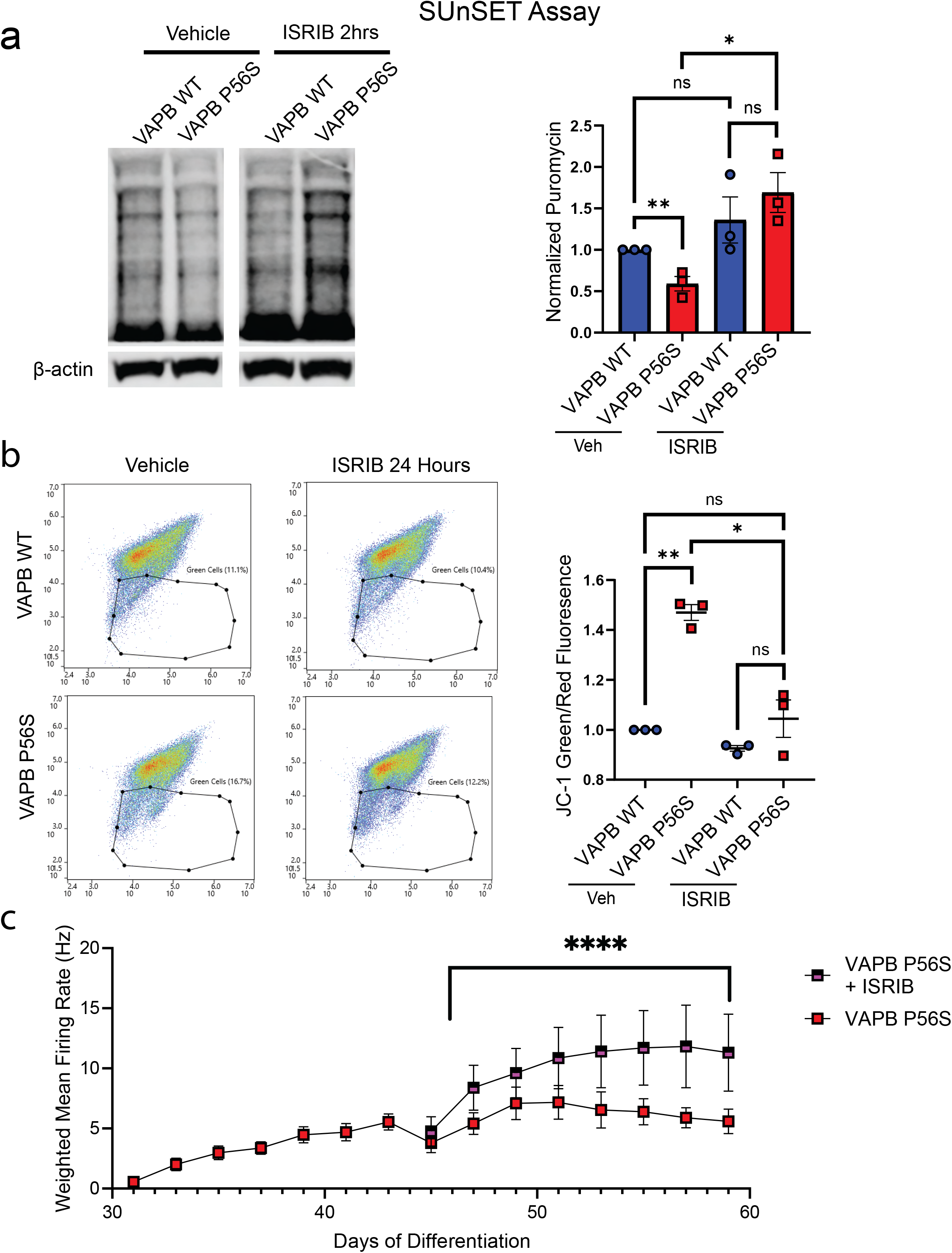
Inhibition of the ISR increases mRNA translation, MMP and neuronal firing in P56S motor neurons. a) SUnSET assay on day 35 of differentiation with vehicle controls and after 2 hours of ISRIB dosing. Mean ± SEM, N=3, unpaired t-test, WT vs P56S *p*=.0024**, P56S vs P56S ISRIB *p*=.0125* b) Scatter plot of day 60 motor neurons stained with JC-1 mitochondrial dye after 24 hours of ISRIB dosing. Mean ± SEM, N=3, paired t-test WT vs P56S *p*=0.0045**, P56S vs P56S ISRIB *p*=.0482*, 2-Way ANOVA for genotype and ISRIB treatment *p*=0.0228* c) Weighted mean firing rate through day 59 of motor neuron differentiation. Mean ± SEM, N=4 wells of an MEA plate for each condition 2-way repeated measure ANOVA from day 45 onward, interaction effect *p*=0.0114*

**Figure 7.**
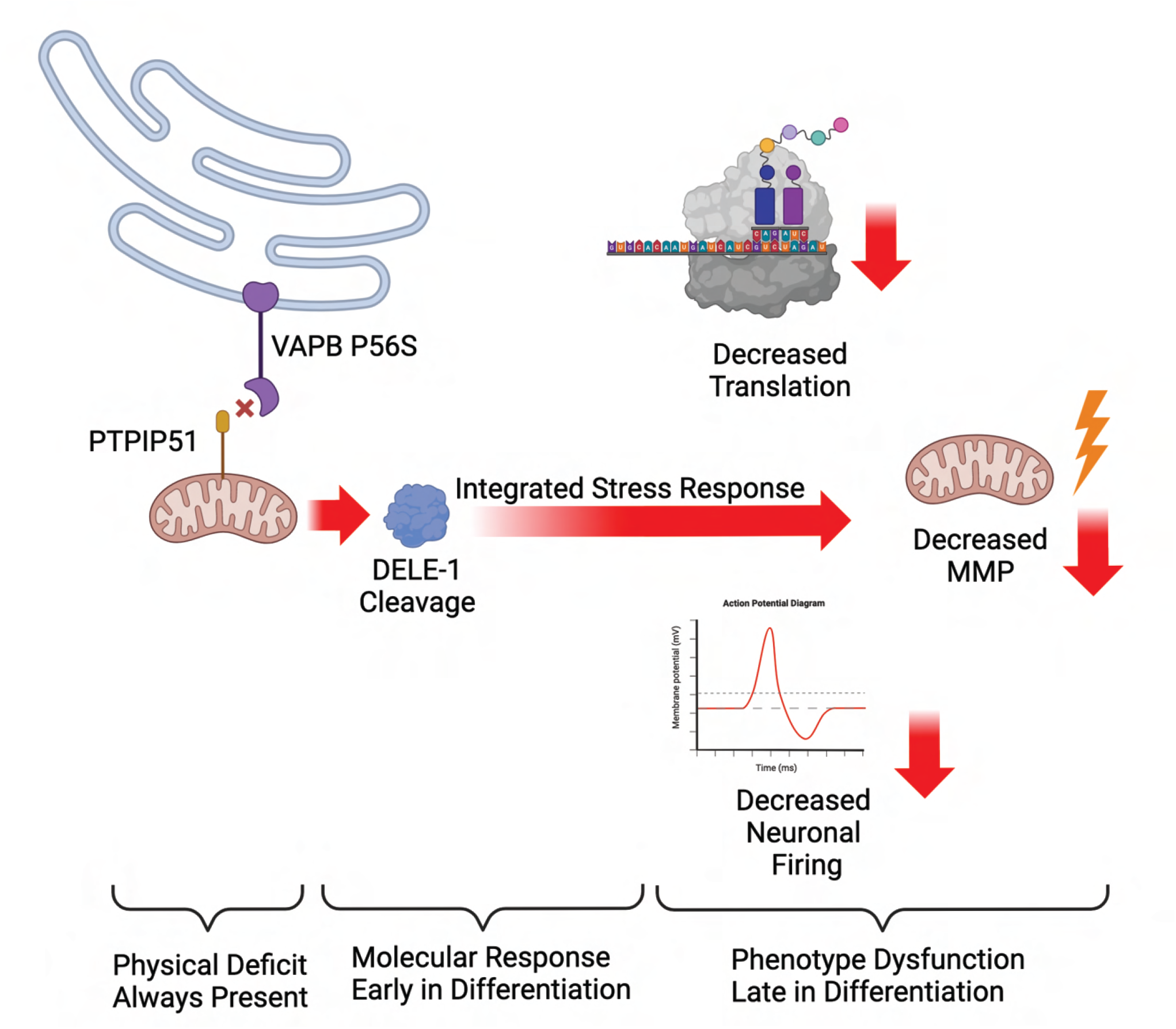
Model of Proposed Pathogenesis. Made with Biorender.com

## DISCUSSION

Lack of full understanding of ALS pathogenesis has hindered effective novel therapy development. Here, we present evidence indicating that the ALS VAPB P56S mutation causes loss of mitochondria-ER contact as well as a sensitization and reduced adaptation to ER stress. Decreased MERC and initiation of the ISR precedes the reduction of MMP and decreased motor neuron electrophysiological activity. Combined with the fact that challenge with stressors such as tunicamycin early in differentiation replicate late-stage disease-relevant phenotype presentation, and that ISRIB reverses these phenotypes, we infer that the underlying physical deficit of decreased MERC initiates ISR activation that potentially intensifies overtime, eventually resulting in the motor neuron degeneration.

Prior investigations have implicated the ISR in ALS pathology, however the actual molecular mechanism associated with variants of familial and sporadic ALS remains elusive. One of the first studies to examine this, showed that ER stress influences disease progression in vulnerable motor neuron populations in SOD1 mice, providing evidence for the link between ER dysfunction and MN degeneration.^32^ Since then, a study showed ISRIB rescuing ER stress caused by SOD1 in culture, however a more recent study described ISRIB-like compounds have the ability to anticipate disease onset and thus shorten SOD1 mice lifespan.^33-35^ Consequently, this lack of consensus in the literature underscores the need for studies in patient derived-cells. While our results agree with other *in vitro* models that eIF2B activators such as ISRIB ameliorate ALS phenotypes, we also acknowledge that their effect *in vivo* require more investigation into the mechanism of action and side effects, given the reported data on aggravation and acceleration of disease progression.^33-35^ Moreover, our results elucidate the biology underlying the defect allowing for alternative therapeutic approaches to be developed targeting these dysfunctional pathways.

Our data support that either decreasing ISR or increasing stress adaptation mechanisms can benefit inhibition of MN-disease in a time and condition dependent manner. To our knowledge this is the most complete pathogenesis of VAPB P56S to date, clearly elucidating the ISR’s role and its link to the mutation. Moreover, we found ISRIB treatment rescues disease-relevant phenotypes caused by the VAPB P56S mutation, including the most clinically relevant, motor neuron electrophysiology. Thus, we suggest that dampening the ISR in motor neurons may be a valid therapeutic approach; interestingly, there is an ongoing Phase I clinical trial investigating this approach (ClinicalTrials.gov Identifier: NCT04948645). While ISR activation has been previously associated with ALS, its identification as the underlying pathogenic mechanism of the VAPB P56S mutation is novel and provides a possible concrete justification for why several studies have seen ISRIB ameliorate ALS phenotypes.^33,35^ There are many forms of ALS, with many similarities between them, pointing to the possibility of a common pathogenesis initiated through disparate means. ^18,33,36^ However, there are multiple concerns about the potential therapeutic application of ISRIB including off-target effects, toxicity, bioavailability, and how effective the treatment would be once motor neuron degeneration has already begun, especially because diagnosis typically occurs after neurodegeneration symptoms present.

While we have elucidated a route for ISR activation in VAPB P56S cells, we have yet to determine the exact mechanism by which ISR activation causes neurodegeneration, another area for possible therapeutic targeting. It could be through the translation suppression from p-eIF2α, either stemming from global translation decrease, or possibly a specific protein affected by the global decrease, more work remains to be done. Identification of the pathophysiological mechanism of disease will allow us to more precisely design drugs to interrupt the disease processes and restore homeostasis to the affected cells. Previous work has shown that the VAPB P56S mutation reduces MERC, however these studies have not proposed a subsequent pathogenesis stemming from this defect.^9,11^ We believe that the detachment of the mitochondria from the ER to be the most likely candidate for the sensitization of the cell to stressors, as it is a clear defect stemming directly from the mutation. This hypothesis has been held before, and even extends to multiple forms of ALS, through a variety of mechanisms, both direct and indirect, such as disruption of the cells redox balance, or as is the case here, direct disruption of the binding of the mitochondria to the ER.^18^ Thus, if this defect shared across ALS types, it will be an important area to focus future efforts on. While ER-mitochondria dissociation has been reported during ALS pathogenesis, here, we present a direct pathway from mutation to phenotype, providing a much clearer path for potential therapeutic development.

In conclusion, we believe that the sensitization and dampened recovery from the Integrated Stress Response to be the causative pathway for the disease progression of ALS Type VIII. While more work remains to be done to link ISR to other forms of ALS, as well as in therapeutic development, we believe these results are a major step forward in elucidating the biological underpinnings of ALS, in pursuit of a viable therapy.

## LIMITATIONS

This study was conducted using only human iPSC-derived motor neurons. Thus, while this work focus on motor neuron-specific molecular mechanisms, we acknowledge that ALS is influenced by many different interactions across multiple cell types. A co-culture or organoids could offer a broader picture of how the VAPB P56S mutation impacts not only the motor neurons directly, but the broader neuronal environment.

## ACKNOWLEDGMENTS

We thank all lab members and CWRU motor neuron group meeting for frequent discussions and valuable input. This work was supported by NINDS K01NS116119, R01NS121374, R61NS133212 to (H.M.), R01NS123524 to (A.E.S.) and DK060596 to (M.H.)

## METHODS

### Antibodies

The following antibodies were used in the experiments described above. Mouse anti-Nanog (SantaCruz Biotechnology; sc-293121), rat anti-SOX2 (ThermoFisher; 14-9811-82), rabbit anti-OCT4 (ThermoFisher; PA5-27438) mouse anti-Nestin (ThermoFisher; MA1-110), chicken anti-GFAP (Abcam; ab4674), rabbit anti-Olig2 (Proteintech; 13999-1-AP), mouse anti-Islet1/2 (DSHB; 39.4D5), rabbit anti-VAPB (MilliporeSigma; SAB1411626 and Proteintech; 14477-1-AP), rabbit anti-PTPI51 (Proteintech; 20641-1-AP), mouse anti-beta-actin (MilliporeSigma; A2228), rabbit anti-ATF4 (Cell Signaling Technologies; 11815), rabbit anti-eIF2alpha (Cell Signaling Technologies; 9722), rabbit anti-phospho-eIF2alpha (EIF2S1 phospho S51) (Abcam; ab32157), mouse anti-puromycin (MilliporeSigma; MABE343), rabbit anti-DELE1 (Proteintech; 21904-1-AP).

### Surface Sensing of Translation Assay

We performed the SUnSET assay based on previously published protocol with some slight modifications.^29^ Briefly, we added puromycin dihydrochloride (Gibco; A11138-03) at 10ug/mL to the motor neuron media and incubated at 37C for 10 minutes. After incubation, we removed the puromycin, washed the cells, and incubated in normal motor neuron media for 45 min at 37C. After this incubation cells were harvested and proteins extracted, with Western Blots being performed as described below, with the caveat that total protein was not quantified and equalized across samples, instead equal volumes of protein extract were loaded, and the samples were normalized through beta actin staining on a separate gel.

### Western Blot

For all the western blots in this paper we harvested cells in Pierce RIPA Lysis and Extraction Buffer (Thermofisher Catalog Number: 89901) (150uL per well of a 6-well plate) with Pierce Protease Inhibitor Tablets (ThermoFisher Catalog Number: A32955) added. Cells were then kept at -80C until ready for protein extraction. Cell pellets were thawed on ice, and then sonicated (Diagenode Bioruptor Pico) for 10 minutes in a 30 second on-off cycle. Solutions were homogenized through repeated pipetting if needed, and then centrifuged at 20,000rcf for 25 minutes at 4C. Supernatant was collected and used to perform a Pierce BCA assay (ThermoFisher Catalog Number: 23227) to quantify total protein. Western blots were run using 20ug of total protein per well on Bolt 4-12% Bis-Tris Plus gels (ThermoFisher Catalog Number: NW04120BOX). Gels were then transferred to Immobilon-P PVDF membrane (Millipore Catalog Number: IPVH00010) using a BioRad TransBlot Turbo transfer station at 2.5A, 25V for 10 minutes. The membrane was then blocked for 1 hour in a 4% milk, 1% Tween-20 in PBS solution on a shaker. The milk buffer was then removed and primary antibodies suspended in LI-COR Intercept blocking buffer (Catalog Number: 927-60001) were added at the dilutions described in the antibody section. Membranes were incubated in primary antibody solution on a shaker overnight at 4C. After incubation, the primary antibody solution was removed and the membrane was rinsed with 1% Tween-20 in PBS (PBS-T) 3 times, then placed on a shaker in PBS-T for 5 minutes. This wash was repeated 3 times and then secondary antibodies (LI-COR IRDye 800CW Donkey anti-Mouse and IRDye 680RD Donkey anti-Rabbit) diluted in 4% milk in PBS-T (1:10,000) were added to the membrane and incubated for 1 hour at room temperature on a shaker. The membrane was then washed as previously in PBS-T and imaged on a LI-COR Odyssey FC imager. Quantification was performed using the Image Studio Lite or Empiria Studio Software.

### JC-1 Assay

The JC-1 assay was performed according to the protocol outlined in the MitoProbe JC-1 Assay Kit (ThermoFisher Catalog Number: M34152) with a few minor modifications. Briefly, motor neurons were dissociated using a 1:1 Accutase:Accumax solution (1mL per well in a 6-well plate), incubated at 37C for 15 minutes. Once the neurons had lifted from the plate, 1mL of Day 25+ motor neuron media was added and the neurons were gently pipetted up and down 1-2 times with a P1000 to homogenize them. The cells were then put through a 70uM mesh strainer, counted, and centrifuged at 200rcf for 2 minutes. Supernatant was removed and the cells were resuspended at a concentration of 1x10^6^ cells/mL in Day 25+ motor neuron media. 1mL of the cells was then placed into a 1.5mL Eppendorf and 8uL of 200uM JC-1 dye was added to each tube (increased to 10uL for day 60 motor neurons). These were then incubated at 37C, 45 minutes for day 35 neurons, and 75 minutes for day 60 neurons as some day 60 motor neurons remained unstained after only 45-minute incubation. After incubation the cells were centrifuged at 200rcf for 90 seconds and the supernatant removed. Cells were then washed with 1mL of sterile PBS. This wash step was repeated twice more with the cells being resuspended in 750uL of sterile PBS filtered through a 0.22uM filter on the final wash. The resulting solution was then run through a Nanocellect WOLF G1 sorter with the gains for FL1, FL2 and FL3 all set to 200mV. Data was compensated with FL1 to FL2 spillover set to 22% and FL2 to FL1 spillover set to 6%. Samples were run until at least 50,000 events were recorded and green fluorescent populations were then gated out by eye, assisted by the population density heat mapping function to ascertain population centers. Percentage of the total population fluorescing green was then normalized to the WT control of each individual biological replicate.

### Proteomics

20uL of Dynabeads Protein A (Fisher Scientific Catalog Number: 100001D) were incubated with 4ug of VAPB antibody overnight. Cells were then harvested with 300uL of Martina lysis buffer (25mM HEPES pH 7.4, 150mM NaCl, 5mM EDTA, 1% Triton X-100) with Pierce Protease Inhibitor Tablets (ThermoFisher Catalog Number: A32955) added. The cells were then homogenized using a 5mL syringe and 25G needle then centrifuged at 20,000 rcf for 25 minutes at 4C. Supernatant was collected and used to perform a Pierce BCA assay (ThermoFisher Catalog Number: 23227) to quantify total protein. A magnetic rack was then used to hold the beads (now conjugated to the antibody) in place while the antibody solution was removed and 1mg of total protein extract was added to the beads. The beads-protein solution was then incubated for 2 hours at 4C on a rotator. Then, using a magnetic rack, the beads were held in place and the supernatant was removed and the beads were washed with lysis buffer three times and submitted to the Proteomics Core at the Lerner Research Institute. The method used for identification was adopted from previous literature.^37^

### Multi-Electrode Array Recordings

Motor neurons were dissociated on day 25 of differentiation and re-plated onto 48-well MEA plate (Axion Biosystems) coated with poly-ornithine (100 ug/mL) and laminin (5 ug/mL), 1x10^5^ cells/well. Media was changed every other day, with recordings being taken with the Maestro Axion Pro for at least 5 minutes, 1 hour post media change, beginning by day 30 of differentiation. Each recording was performed at 37 degrees Celsius and 5% CO2 using the AxIS Software Spontaneous Neural Configuration for spontaneous activity. Recordings were then processed with Axion Biosystems’s Neural Metrics Tool and values for each well were exported and analyzed. Wells with a zero mean firing rate were omitted from analysis as not firing. Bursts were identified using an interspike interval (ISI) threshold requiring a 5-spike minimum and 25-ms maximum ISI.

### Electron Microscopy

Motor neurons were cultured as normal and then roughly 7 days before fixation they were dissociated and replated onto 12 mm Snapwell™ Insert with 0.4 µm Pore Polyester Membranes (Corning Product #3801). They were then fixed at their respective timepoints by removing the growth medium, washing the plate and adding 2-3mL of fixative solution consisting of 2.5% glutaraldehyde in cacodylate buffer (pH 7.3), to the top and bottom of the membrane. The cells were incubated in the fixative solution for 1hr at room temperature. The fixative solution was then removed, and a fresh 2-3mL was once again added to the top and bottom of the membrane and incubated for another hour at room temperature. The fixative solution was removed and then the membrane was washed with PBS 3 times, leaving the PBS on the membrane for 5 minutes each time. The wells with membrane inserts were then completely filled with PBS and kept at 4C until they were transported to the Cryo-Electron Microscopy Core Facility at the Cleveland Center for Membrane and Structural Biology for processing and imaging. The specimen was postfixed in ferrocyanide-reduced 1% osmium tetroxide.^38^ After a soak in acidified uranyl acetate, the specimen was dehydrated in ethanol, passed through propylene oxide, and embedded in Embed-812 (Electron Microscopy Science, PA).^39^ Sections were cut in a horizontal plane parallel to that of the membrane to provide panoramic views of the cells. Thin sections were stained first with acidified uranyl acetate in 50% methanol, then with the triple lead stain of Sato as modified by Hanaichi et al.^39,40^ These sections were examined in a FEI Tecnai Spirit (T12) with a Gatan US4000 4kx4k CCD.

### Tunicamycin

Tunicamycin (Tocris Catalog #3516) was resuspended in DMSO at a concentration of 5mg/mL. When dosing cells it was added to the medium at a final working concentration of 7.5ug/mL, with this concentration being used for all results shown here.

### ISRIB

ISRIB (Sigma-Millipore Catalog #SML0843) was resuspended in DMSO at a concentration of 5mg/mL. When dosing cells it was added to the medium at a final working concentration of 5ug/mL, with the exception of the neuronal firing rescue experiment (Figure 6C) for which it was added for a final concentration of 15ug/mL. ISRIB was added every 8 hours for the JC-1 ISRIB rescue experiment (Figure 6B) and every 24 hours for MEA recordings (Figure 6C).

## Notes

### Competing Interest Statement

The authors have declared no competing interest.

## REFERENCE

1 Hardiman, O., van den Berg, L. H. & Kiernan, M. C. Clinical diagnosis and management of amyotrophic lateral sclerosis. Nature Reviews Neurology 7, 639–649, doi:10.1038/nrneurol.2011.153 (2011).

2 Nishimura, A. L. et al. A Mutation in the Vesicle-Trafficking Protein VAPB Causes Late-Onset Spinal Muscular Atrophy and Amyotrophic Lateral Sclerosis. The American Journal of Human Genetics 75, 822–831, doi:10.1086/425287 (2004).

3 Mitne-Neto, M. et al. A mutation in human VAP-B--MSP domain, present in ALS patients, affects the interaction with other cellular proteins. Protein Expr Purif 55, 139–146, doi:10.1016/j.pep.2007.04.007 (2007).

4 Borgese, N., Navone, F., Nukina, N. & Yamanaka, T. Mutant VAPB: Culprit or Innocent Bystander of Amyotrophic Lateral Sclerosis? Contact 4, 25152564211022515, doi:10.1177/25152564211022515 (2021).

5 Borgese, N., Iacomino, N., Colombo, S. F. & Navone, F. The Link between VAPB Loss of Function and Amyotrophic Lateral Sclerosis. Cells 10, doi:10.3390/cells10081865 (2021).

6 Kirmiz, M., Vierra, N. C., Palacio, S. & Trimmer, J. S. Identification of VAPA and VAPB as Kv2 Channel-Interacting Proteins Defining Endoplasmic Reticulum-Plasma Membrane Junctions in Mammalian Brain Neurons. J Neurosci 38, 7562–7584, doi:10.1523/jneurosci.0893-18.2018 (2018).

7 Lev, S., Ben Halevy, D., Peretti, D. & Dahan, N. The VAP protein family: from cellular functions to motor neuron disease. Trends Cell Biol 18, 282–290, doi:10.1016/j.tcb.2008.03.006 (2008).

8 Mitne-Neto, M. et al. Downregulation of VAPB expression in motor neurons derived from induced pluripotent stem cells of ALS8 patients. Hum Mol Genet 20, 3642–3652, doi:10.1093/hmg/ddr284 (2011).

9 Stoica, R. et al. ER–mitochondria associations are regulated by the VAPB–PTPIP51 interaction and are disrupted by ALS/FTD-associated TDP-43. Nat Commun 5, 3996, doi:10.1038/ncomms4996 (2014).

10 Obara, C. J. et al. Motion of VAPB molecules reveals ER–mitochondria contact site subdomains. Nature, doi:10.1038/s41586-023-06956-y (2024).

11 Gómez-Suaga, P. et al. The VAPB-PTPIP51 endoplasmic reticulum-mitochondria tethering proteins are present in neuronal synapses and regulate synaptic activity. Acta Neuropathologica Communications 7, 35, doi:10.1186/s40478-019-0688-4 (2019).

12 Teuling, E. et al. Motor neuron disease-associated mutant vesicle-associated membrane protein-associated protein (VAP) B recruits wild-type VAPs into endoplasmic reticulum-derived tubular aggregates. J Neurosci 27, 9801–9815, doi:10.1523/jneurosci.2661-07.2007 (2007).

13 Kanekura, K., Nishimoto, I., Aiso, S. & Matsuoka, M. Characterization of amyotrophic lateral sclerosis-linked P56S mutation of vesicle-associated membrane protein-associated protein B (VAPB/ALS8). J Biol Chem 281, 30223–30233, doi:10.1074/jbc.M605049200 (2006).

14 Cabukusta, B. et al. Human VAPome Analysis Reveals MOSPD1 and MOSPD3 as Membrane Contact Site Proteins Interacting with FFAT-Related FFNT Motifs. Cell Rep 33, 108475, doi:10.1016/j.celrep.2020.108475 (2020).

15 James, C. et al. Proteomic mapping by rapamycin-dependent targeting of APEX2 identifies binding partners of VAPB at the inner nuclear membrane. Journal of Biological Chemistry 294, 16241–16254, doi:10.1074/jbc.RA118.007283 (2019).

16 Gomez-Suaga, P. et al. The ER-Mitochondria Tethering Complex VAPB-PTPIP51 Regulates Autophagy. Curr Biol 27, 371–385, doi:10.1016/j.cub.2016.12.038 (2017).

17 Markmiller, S. et al. Context-Dependent and Disease-Specific Diversity in Protein Interactions within Stress Granules. Cell 172, 590–604.e513, doi:10.1016/j.cell.2017.12.032 (2018).

18 Chen, J., Bassot, A., Giuliani, F. & Simmen, T. Amyotrophic Lateral Sclerosis (ALS): Stressed by Dysfunctional Mitochondria-Endoplasmic Reticulum Contacts (MERCs). Cells 10, doi:10.3390/cells10071789 (2021).

19 Pakos-Zebrucka, K. et al. The integrated stress response. EMBO Rep 17, 1374–1395, doi:10.15252/embr.201642195 (2016).

20 B’chir, W. et al. The eIF2α/ATF4 pathway is essential for stress-induced autophagy gene expression. Nucleic Acids Research 41, 7683–7699, doi:10.1093/nar/gkt563 (2013).

21 Kilberg, M. S., Shan, J. & Su, N. ATF4-dependent transcription mediates signaling of amino acid limitation. Trends in Endocrinology & Metabolism 20, 436–443, doi:10.1016/j.tem.2009.05.008 (2009).

22 Fessler, E. et al. A pathway coordinated by DELE1 relays mitochondrial stress to the cytosol. Nature 579, 433–437, doi:10.1038/s41586-020-2076-4 (2020).

23 Bauer, B. N., Rafie-Kolpin, M., Lu, L., Han, A. & Chen, J. J. Multiple autophosphorylation is essential for the formation of the active and stable homodimer of heme-regulated eIF2alpha kinase. Biochemistry 40, 11543–11551, doi:10.1021/bi010983s (2001).

24 Rafie-Kolpin, M., Han, A. P. & Chen, J. J. Autophosphorylation of threonine 485 in the activation loop is essential for attaining eIF2alpha kinase activity of HRI. Biochemistry 42, 6536–6544, doi:10.1021/bi034005v (2003).

25 Costa-Mattioli, M. & Walter, P. The integrated stress response: From mechanism to disease. Science 368, doi:10.1126/science.aat5314 (2020).

26 Mick, E. et al. Distinct mitochondrial defects trigger the integrated stress response depending on the metabolic state of the cell. eLife 9, e49178, doi:10.7554/eLife.49178 (2020).

27 Baird, T. D. & Wek, R. C. Eukaryotic initiation factor 2 phosphorylation and translational control in metabolism. Adv Nutr 3, 307–321, doi:10.3945/an.112.002113 (2012).

28 Harding, H. P. et al. Regulated translation initiation controls stress-induced gene expression in mammalian cells. Mol Cell 6, 1099–1108, doi:10.1016/s1097-2765(00)00108-8 (2000).

29 Schmidt, E. K., Clavarino, G., Ceppi, M. & Pierre, P. SUnSET, a nonradioactive method to monitor protein synthesis. Nat Methods 6, 275–277, doi:10.1038/nmeth.1314 (2009).

30 Goodman, C. A. & Hornberger, T. A. Measuring protein synthesis with SUnSET: a valid alternative to traditional techniques? Exerc Sport Sci Rev 41, 107–115, doi:10.1097/JES.0b013e3182798a95 (2013).

31 Zyryanova, A. F. et al. ISRIB Blunts the Integrated Stress Response by Allosterically Antagonising the Inhibitory Effect of Phosphorylated eIF2 on eIF2B. Mol Cell 81, 88–103.e106, doi:10.1016/j.molcel.2020.10.031 (2021).

32 Saxena, S., Cabuy, E. & Caroni, P. A role for motoneuron subtype–selective ER stress in disease manifestations of FALS mice. Nature Neuroscience 12, 627–636, doi:10.1038/nn.2297 (2009).

33 Marlin, E., Viu-Idocin, C., Arrasate, M. & Aragón, T. The Role and Therapeutic Potential of the Integrated Stress Response in Amyotrophic Lateral Sclerosis. Int J Mol Sci 23, doi:10.3390/ijms23147823 (2022).

34 Marlin, E. et al. Pharmacological inhibition of the integrated stress response accelerates disease progression in an amyotrophic lateral sclerosis mouse model. British Journal of Pharmacology n/a, doi:10.1111/bph.16260.

35 Bugallo, R. et al. Fine tuning of the unfolded protein response by ISRIB improves neuronal survival in a model of amyotrophic lateral sclerosis. Cell Death & Disease 11, 397, doi:10.1038/s41419-020-2601-2 (2020).

36 Mejzini, R. et al. ALS Genetics, Mechanisms, and Therapeutics: Where Are We Now? Frontiers in neuroscience 13, 1310–1310, doi:10.3389/fnins.2019.01310 (2019).

37 Mohammed, H. et al. Rapid immunoprecipitation mass spectrometry of endogenous proteins (RIME) for analysis of chromatin complexes. Nature Protocols 11, 316–326, doi:10.1038/nprot.2016.020 (2016).

38 Karnovsky, M. in Proc. 11th Meeting Am. Soc. Cell Biol. New Orleans, LA (abstract).

39 Tandler, B. Improved uranyl acetate staining for electron microscopy. J Electron Microsc Tech 16, 81–82, doi:10.1002/jemt.1060160110 (1990).

40 Hanaichi, T. et al. A stable lead by modification of Sato’s method. J Electron Microsc (Tokyo) 35, 304–306 (1986).

