## Supplemental Figures for "Mitochondrial dysfunction heightens the integrated stress response to drive ALS pathogenesis"

Extended Data Figure 1: Characterization of VAPB doxycycline-inducible iPSC. a) Next-generation sequencing of the ALS iPSC after CRSIPR-Cas9 mediated genome editing. Bi-allelic frameshift mutations were identified. b) Immunofluorescence of the doxycycline-inducible iPSC lines, each line showing full expression of pluripotency markers Nanog, SOX2, and Oct4. N=3 replicates, a minimum of 3 images per differentiation. d) Quantitative PCR of VAPB HA (to only quantify VAPB transcripts from the constructs) on the doxycycline-inducible lines as iPSCs. GAPDH and RPL3 were used for normalization. Mean ± SEM, N=3 qPCR replicates per line, per time point.

Extended Data Figure 2: Characterization of VAPB doxycycline-inducible motor neurons. a) Western blot for VAPB on days 15 (motor neuron progenitors, MNP) and 30 of motor neuron differentiation. Unaffected and Affected are familial patients iPSC, with the doxycycline-inducible lines made from the Affected line. Mean ± SEM, N=3 b) Immunofluorescence of the doxycycline-inducible lines on day 15 of motor neuron differentiation, each line showing full expression of motor neuron progenitor markers Nestin and Olig2, with no expression of the glial marker GFAP. N=3 replicates, minimum of 3 images per differentiation. c) Immunofluorescence of the doxycycline-inducible lines on day 30 of motor neuron differentiation, comparing the number of double positive DAPI and Islet 1/2 cells. *p*-value>0.05, Mean ± SEM, N=3 replicates, minimum of 3 images per differentiation.

Extended Data Figure 3: High Exposure of VAPB Coimmunoprecipitation western blot.

Extended Data Figure 4: Mitochondria-ER Contact Sites. a) Number of mitochondria counted for each line across all images taken at that time point. N=1. b) Percent of mitochondria present in each image with no ER-mitochondrial contact observable. *p*-values: VAPB WT Day 35 vs VAPB P56S Day 35=0.0248*, VAPB WT Day 35 vs VAPB WT Day 60=0.0078**, VAPB WT Day 35 vs VAPB P56S Day 60=0.0293*, Violin plot with median, first and third quartiles denoted as dashed lines, N=1 replicate submitted for EM, 13-21 images analyzed for each condition. c) Area of mitochondria in pixels, all *p*-values>0.05. Violin plot with median, first, and third quartiles denoted as dashed lines, from sample submitted for EM, N=13-21 images analyzed for each condition.

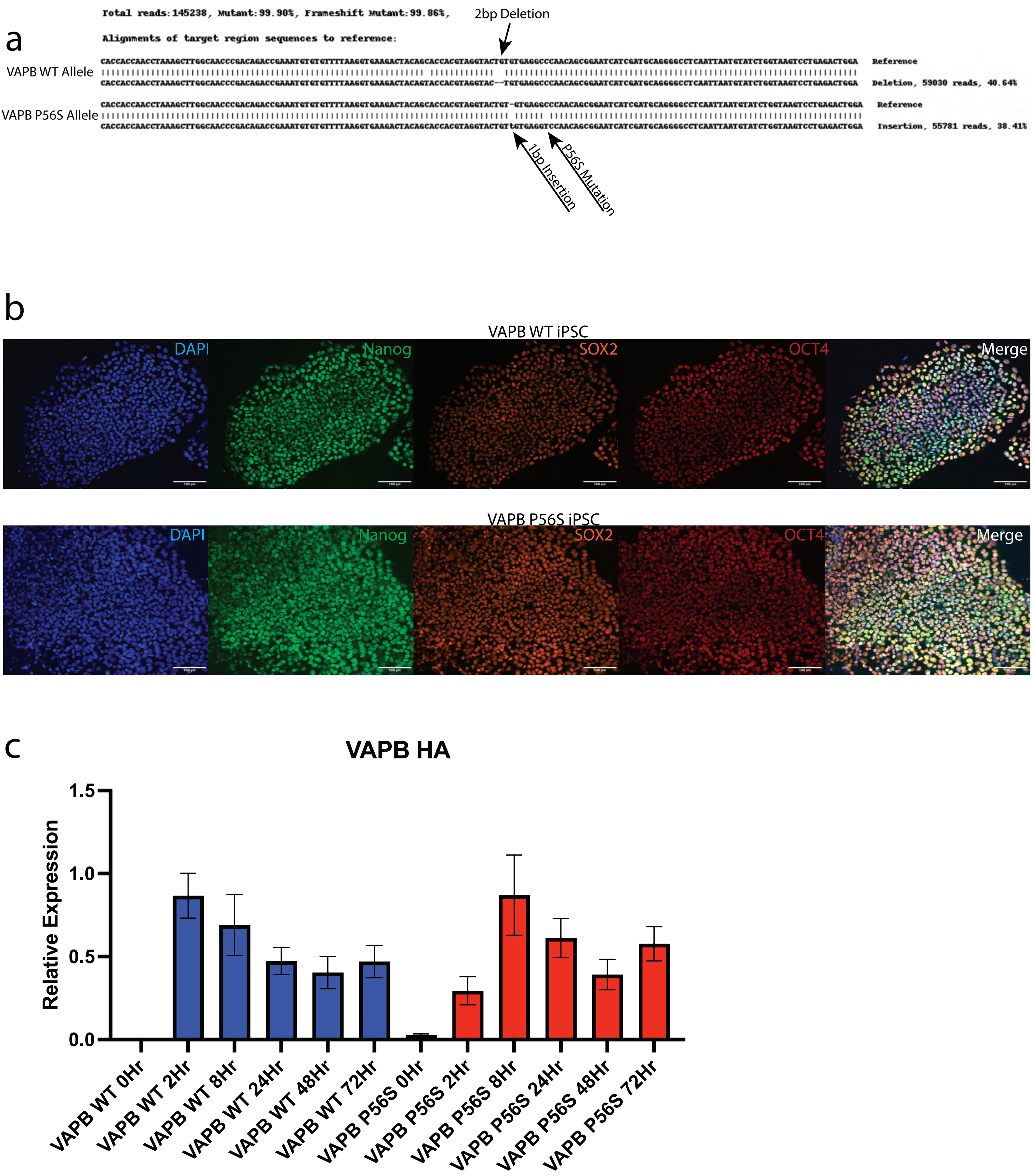
a
Total reads:145238, Mutant:99.90%, Frameshift Mutant:99.86%,
Alignments of target region sequences to reference:
2bp Deletion
VAPB WT Allele
Reference
Deletion, 59030 reads, 40.64%
VAPB P56S Allele
Reference
Insertion, 55781 reads, 38.41%
1bp Insertion
P56S Mutation
b
VAPB WT iPSC
DAPI
Nanog
SOX2
OCT4
Merge
VAPB P56S iPSC
DAPI
Nanog
SOX2
OCT4
Merge
c
VAPB HA
Relative Expression
1.5
1.0
0.5
0.0
VAPB WT 0Hr
VAPB WT 2Hr
VAPB WT 8Hr
VAPB WT 24Hr
VAPB WT 48Hr
VAPB WT 72Hr
VAPB P56S 0Hr
VAPB P56S 2Hr
VAPB P56S 8Hr
VAPB P56S 24Hr
VAPB P56S 48Hr
VAPB P56S 72Hr

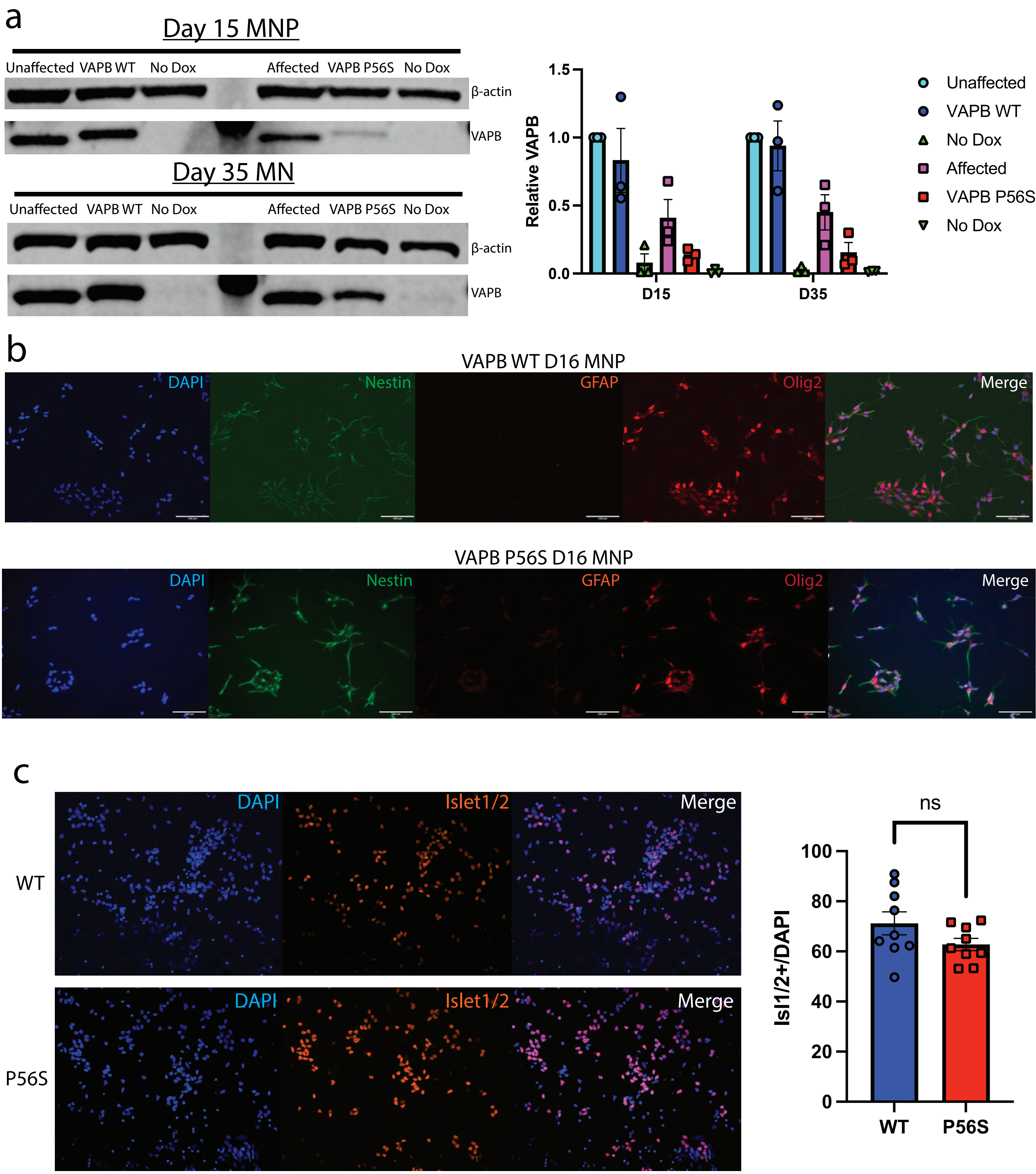
a
Day 15 MNP
Unaffected VAPB WT No Dox Affected VAPB P56S No Dox
β-actin
VAPB
Day 35 MN
Unaffected VAPB WT No Dox Affected VAPB P56S No Dox
β-actin
VAPB
Relative VAPB
1.5
1.0
0.5
0.0
D15
D35
Unaffected
VAPB WT
No Dox
Affected
VAPB P56S
No Dox
b
VAPB WT D16 MNP
DAPI
Nestin
GFAP
Olig2
Merge
VAPB P56S D16 MNP
DAPI
Nestin
GFAP
Olig2
Merge
c
WT
P56S
DAPI
Islet1/2
Merge
ns
Isl1/2+/DAPI
100
80
60
40
20
0
WT
P56S

a

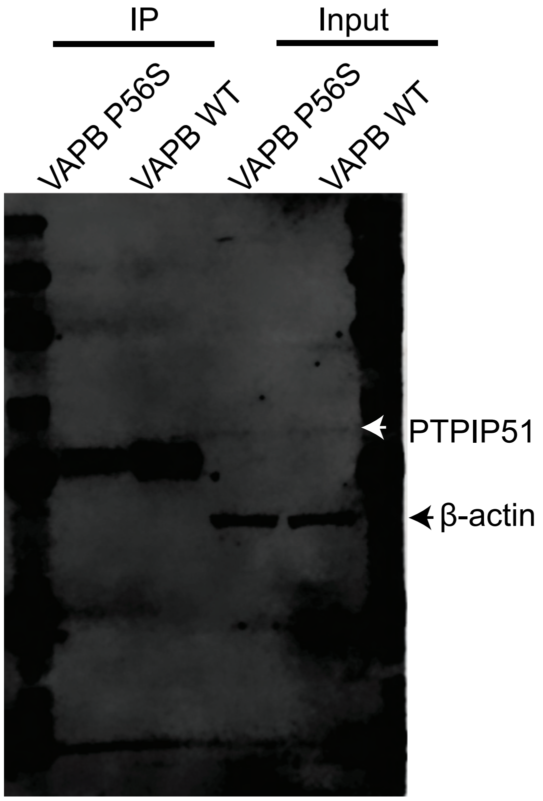

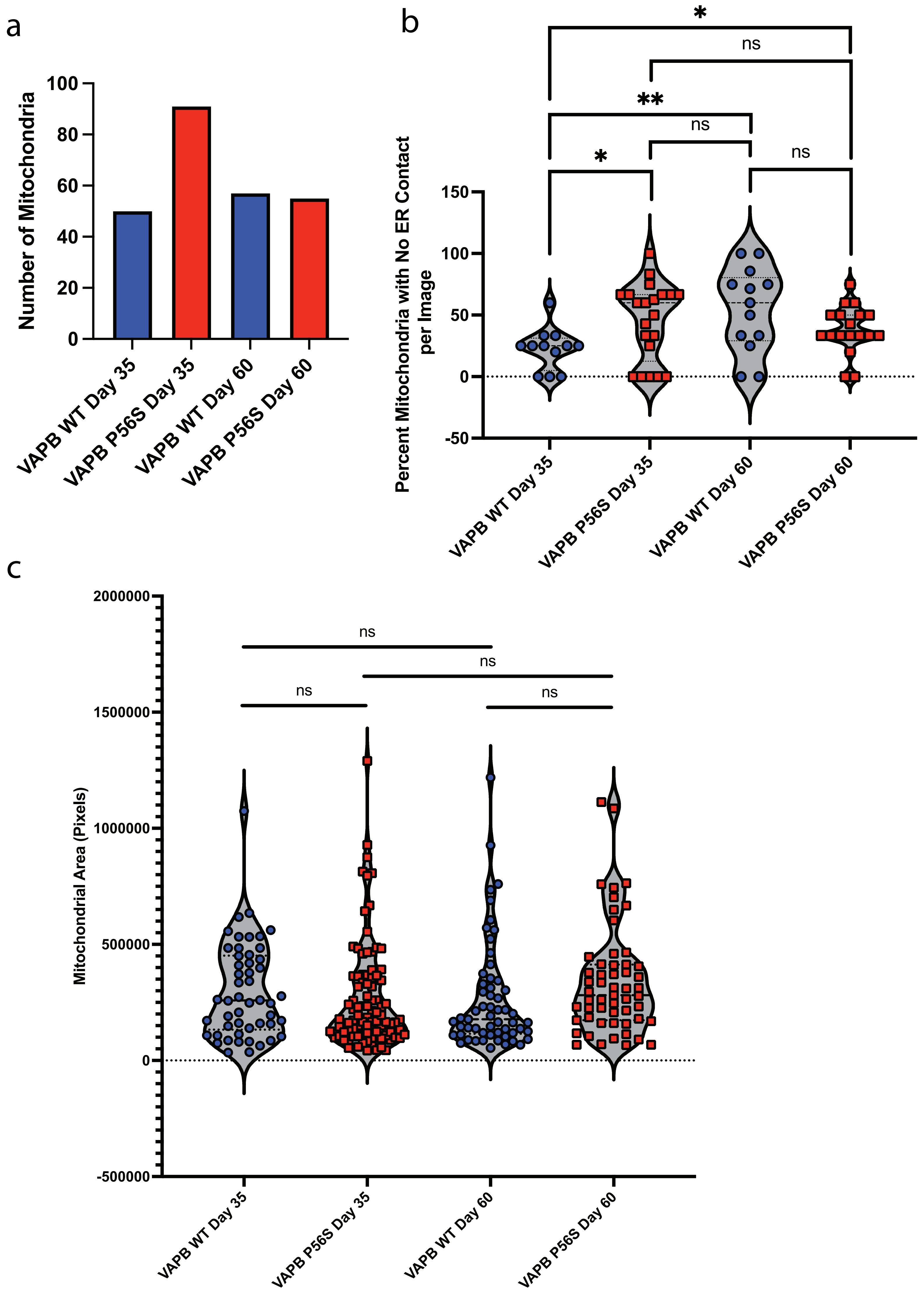

a
Number of Mitochondria
100
80
60
40
20
0
VAPB WT Day 35
VAPB P56S Day 35
VAPB WT Day 60
VAPB P56S Day 60
b
Percent Mitochondria with No ER Contact per Image
150
100
50
0
-50
*
ns
**
ns
*
ns
VAPB WT Day 35
VAPB P56S Day 35
VAPB WT Day 60
VAPB P56S Day 60
c
Mitochondrial Area (Pixels)
2000000
1500000
1000000
500000
0
-500000
ns
ns
ns
ns
VAPB WT Day 35
VAPB P56S Day 35
VAPB WT Day 60
VAPB P56S Day 60
